# Facilitation-inhibition control of motor neuronal persistent inward currents in young and older adults

**DOI:** 10.1101/2022.08.08.503135

**Authors:** Lucas B. R. Orssatto, Gabriel L. Fernandes, Anthony J. Blazevich, Gabriel S. Trajano

## Abstract

A well-coordinated control of motor neuronal persistent inward currents (PICs) via diffuse neuromodulation and local inhibition is essential to ensure motor units discharge at required times and frequencies. Current best estimates indicate that PICs are reduced in older adults; however, it is not yet known whether PIC facilitation-inhibition control is also altered with ageing. We investigated the responses of PICs to i) a remote handgrip contraction, which is believed to diffusely increase serotonergic input onto motor neurones, and ii) tendon vibration of the antagonist muscle, which elicits reciprocal inhibition, in both young and older adults. High-density surface electromyograms were collected from *soleus* and *tibialis anterior* of 18 young and 26 older adults during triangular-shaped plantar and dorsiflexion contractions to 20% (handgrip experiments) and 30% (vibration experiments) of maximum torque (rise-decline rate of 2%/s). The paired-motor-unit analysis was used to calculate ΔF, which is assumed proportional to PIC strength. ΔF increased in both *soleus* (0.55pps, 16.0%) and *tibialis anterior* (0.42pps, 11.4%) during the handgrip contraction independent of age. However, although antagonist tendon vibration reduced ΔF in *soleus* (0.28pps, 12.6%) independent of age, less reduction was observed in older (0.42pps, 10.7%) than young adults (0.72pps, 17.8%) in *tibialis anterior*. Our data indicate a preserved ability of older adults to amplify PICs following a remote handgrip contraction, during which increased serotonergic input onto the motor neurones is expected, in both lower leg muscles. However, PIC deactivation in response to reciprocal inhibition was impaired with ageing in *tibialis anterior* despite being preserved in *soleus*.

**KEYPOINTS:**

- Motor neuronal persistent inward currents (PICs) are amplified via diffuse neuromodulation and deactivated by local inhibition to ensure motor units discharge at required times and frequencies, allowing a normal motor behaviour.
- PIC amplitudes appear to be reduced with ageing, however it is not known whether PIC facilitation-inhibition control is also altered.
- Remote handgrip contraction, which should diffusely increase serotonergic input onto motor neurones, amplified PICs similarly in both *soleus* and *tibialis anterior* of young and older adults.
- Antagonist tendon vibration, which induces reciprocal inhibition, reduced PICs in *soleus* in both young and older adults but had less effect in *tibialis anterior* in older adults.
- Our data suggest that older adults have preserved *soleus* PIC facilitation during low-intensity contractions, equivalent to activities such as standing and walking. However, a reduced reciprocal inhibition of PICs in *tibialis anterior* may contribute to locomotion impairments, such as increases in *soleus-tibialis anterior* co-activation during propulsion.

## INTRODUCTION

Persistent inward currents (PICs), generated by voltage-gated sodium and calcium channels within the motor neurones, enable amplification and prolongation of excitatory synaptic input (Heckman *et al*., 2005). Motor neurones can adjust PIC strength via a fine control between neuromodulation and inhibition according to the desired task demands (Heckman *et al*., 2005). This neuromodulatory control of PICs is dictated by the amount of serotonin and noradrenaline released from the brainstem nuclei and delivered onto the motor neurone dendrites, activating slow activating L-type Ca^2+^ and fast activating persistent Na^+^ currents (Heckman *et al*., 2008). It has also been suggested that monoaminergic projections to the spinal cord vary their activity proportional to the demand (e.g., intensity) of the performed motor activity. Thus, the stronger PIC amplification observed in higher intensity activities (Orssatto *et al*., 2021*b*) could theoretically result from a higher monoaminergic input onto the motor neurones (Lee & Heckman, 1999, 2000; Heckman *et al*., 2005). However, the descending monoaminergic projections are highly diffuse, resulting in increased excitation of diverse muscle groups, including those not involved in the desired tasks (Heckman *et al*., 2008; Wei *et al*., 2014). For example, contraction of one muscle group triggers a serotonergic-mediated increase in motor neuronal gain in other, unrelated muscles (Wei *et al*., 2014). Alternatively, deactivation of undesired PICs in specific motor neurones can be achieved by local inhibitory circuits in the spinal cord. PICs are highly sensitive to inhibition and are thus turned off by inhibitory input (Hounsgaard *et al*., 1988; Hultborn *et al*., 2003; Kuo *et al*., 2003; Revill & Fuglevand, 2017; Mesquita *et al*., 2022; Pearcey *et al*., 2022). For example, reciprocal inhibition can deactivate PICs from the antagonist muscles (Hyngstrom *et al*., 2007; Vandenberk & Kalmar, 2014; Mesquita *et al*., 2022; Pearcey *et al*., 2022), avoiding undesirable coactivations. Thus, a well-coordinated control of PICs via diffuse activation and local deactivation is essential to ensure motor units discharge at desired times and frequencies, allowing normal motor behaviour (Heckman *et al*., 2008).

Age-related alterations within the nervous system (Manini *et al*., 2013; Orssatto *et al*., 2018) contribute to the impairments in movement control and force production observed in older individuals (Suetta *et al*., 2019). Recent estimates suggest that reduced motor neuronal PIC strength plays a role in these impairments (Hassan *et al*., 2021; Orssatto *et al*., 2021*a*). However, it is not known whether PIC neuromodulation-inhibition control is also altered with ageing. Older adults present dysfunctions within the monoaminergic system and inhibitory circuits that could contribute to both inefficient monoaminergic neuromodulation and localised effects of inhibition on PICs (Johnson *et al*., 1993; Ko *et al*., 1997; Míguez *et al*., 999; Kido *et al*., 2004; Hortobágyi *et al*., 2006; Shibata *et al*., 2006; Liu *et al*., 2020; Steinbusch *et al*., 2021), hence potentially contributing to impairments in motor control in this population. Detrimental effects of ageing on the monoaminergic system as well as chronic inflammation reduce serotonin and noradrenaline secretions and thus input onto the motoneurons (Johnson *et al*., 1993; Ko *et al*., 1997; Míguez *et al*., 1999; Shibata *et al*., 2006; Liu *et al*., 2020; Steinbusch *et al*., 2021), which would impair neuromodulation and attenuate PIC amplification. With respect to inhibitory circuits, older adults have reduced cortical and spinal reciprocal inhibition (Kido *et al*., 2004; Hortobágyi *et al*., 2006), which could impair PIC inhibition, generating undesired muscle contractions such as the observed age-related increases in antagonist coactivation (Macaluso *et al*., 2002; Hortobágyi & Devita, 2006; Hortobágyi *et al*., 2009)). Exploring the dynamics of PIC neuromodulation and inhibition in older adults could provide therefore important insight into factors underpinning impaired movement control (e.g., locomotion) in this population.

In the present study, the responses of PICs to i) a remote handgrip contraction, which is believed to diffusely increase serotonergic input onto motor neurones, and ii) reciprocal inhibition, which induces reciprocal inhibition, were estimated in *soleus* and *tibialis anterior* of young and older adults using the gold-standard paired motor unit technique (Gorassini *et al*., 2002; Vandenberk & Kalmar, 2014). To answer these questions, the study involved two experiments. In Experiment 1 we estimated PICs after a remote contraction with upper limb muscles not involved in the *soleus-* and *tibialis anterior-*targeting tasks, theoretically inducing increases in serotonergic input onto the motor neurones by taking advantage of their diffuse descending monoaminergic projections into the spinal cord (Heckman *et al*., 2008; Wei *et al*., 2014). In Experiment 2, we estimated PIC strength while activating a well-known disynaptic reciprocal inhibition circuit using tendon vibration of the antagonist muscle (Pearcey *et al*., 2022), which activates Ia inhibitory inter neurones via antagonist Ia afferent stimulation by stimulating its muscle spindles (Burke *et al*., 1976; Grande & Cafarelli, 2003; Pearcey *et al*., 2022). We hypothesised that i) older adults would present smaller PIC increases than young adults in response to a remote handgrip contraction, and ii) older adults would present reduced attenuation of PICs in response to reciprocal inhibition, irrespective of the tested muscles. *Soleus* and *tibialis anterior* were chosen in this study due to their important antagonist co-involvement in postural stability and locomotion (Polcyn *et al*., 1998; Laughton *et al*., 2003; Hortobágyi & Devita, 2006).

## METHODS

### Participants and Ethical Procedures

Eighteen young adults aged 18-35 years and 26 non-sarcopenic older adults aged ≥ 65 years volunteered to the present study. They had no history of neurological disorders, were free of lower limb musculoskeletal injuries, and were not taking medications that could influence the monoaminergic system, including serotonin or noradrenaline modulators (e.g., beta-blockers and serotonin reuptake inhibitors). They were instructed to not consume caffeinated foods or beverages 24 h before the testing session. One young and one older adult were excluded from the study because no motor units were identified in *soleus* and *tibialis anterior*. The participants had already completed a previous study in our laboratory from other tests conducted within the same visit, before the procedures of the present study were imposed. Further participant characteristics and other variables can be found in that publication (Orssatto *et al*., 2021*a*). This study was approved by the University Human Research Ethics Committee, and all participants gave written informed consent before participating. Data collection was conducted during the COVID-19 pandemic and all safety procedures followed the local state government policies.

### Study design and testing procedures

Participants visited the laboratory on a single occasion. Initially, participants performed a warm-up and three plantar flexion, dorsiflexion, and handgrip maximal voluntary isometric contractions (MVC, ∼3-s with 30-s rest intervals) and were familiarised to the submaximal ramped contractions. Subsequently, they performed four submaximal ramp-shaped contractions, which have been already analysed and the data published (Orssatto *et al*., 2021*a*). After 10 min of rest, the procedures from the present study were conducted. The present cross-sectional study was divided into two experiments testing plantar and dorsiflexion tasks, which are described below. In Experiments 1 and 2, participants were seated upright in the chair of an isokinetic dynamometer (Biodex System 4, Biodex Medical system, Shirley, NY) with the knee fully extended (0°) and ankle in anatomical position (0°).

#### Experiment 1

Participants performed two sets of two ramp-shaped contractions to 20% of their peak torque (10-s up and 10-s down), interspersed by an ipsilateral remote handgrip contraction or resting control condition (Figure 1A). Contracting a muscle unrelated to the tested task is a non-invasive method to induce serotonergic-mediated gain on the tested muscle (Wei *et al*.,2014). The remote handgrip contraction was preceded by a 10-s preparation period, followed by a 30-s contraction to 40% of their maximal handgrip force. Participants were asked to avoid any movement (i.e., muscle contraction) during the resting control condition, which lasted 40 s. The subsequent ramp-shaped contractions were performed immediately after the remote handgrip or resting control conditions (i.e., remote contraction was ceased before plantar- or dorsiflexion contractions commenced so dual tasking was avoided). The order in which each condition was performed and which each muscle was tested were randomised. Participants received real-time visual torque-trace feedback during each contraction and were instructed to follow the torque path displayed on a 58-cm computer monitor. When an abrupt increase or decrease in torque was observed (i.e., the torque trajectory was not closely followed), the whole trial was excluded and repeated 2 min later.

**Figure 1.**
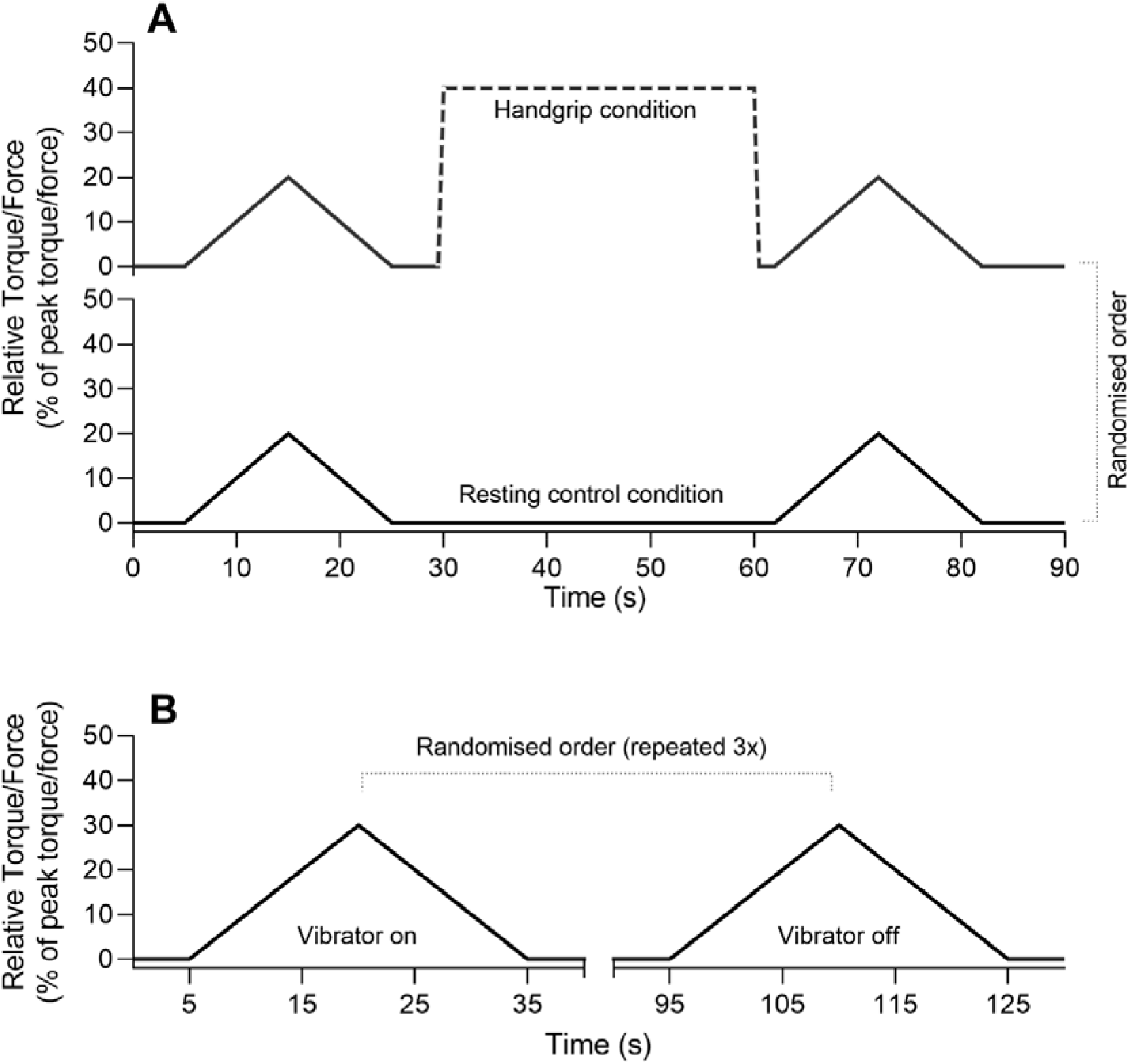
Design of Experiments 1 and 2. Panel A presents Experiment 1 design with the handgrip condition shown at top and control condition at bottom. Panel B shows Experiment 2 design.

#### Experiment 2

Participants performed six ramp-shaped contractions to 30% of their peak torque at a rate of torque rise and decline of 2%/s (15-s up and 15-s down). Three contractions were performed with high-frequency vibration applied to the tendon of the antagonist muscles and three with the vibrator device turned off (i.e., control condition). Conditions were alternated in a randomised order, interspersed by 1-min rest intervals (Figure 1B). Tendon vibration (115 Hz, Vibrasens Proprioceptive Technology, France) was applied to the Achilles tendon during the dorsiflexion contractions and to the *tibialis anterior* tendon during the plantar flexion contractions. Vibration commenced 10 s before the start of the ramp-shaped contraction and continued until 2 s after the torque trace returned to baseline, and was thus imposed through the duration of the ramp contraction.

### Surface electromyography

Skin preparation included shaving, abrasion, and cleansing each site with 70% isopropyl alcohol. Two semi-disposable 32-channel electrode grids with a 10-mm interelectrode distance (ELSCH032NM6, OTBioelettronica, Torino, Italy) were placed over the medial and lateral portions of *soleus* (either side of the Achilles tendon) and another two electrode grids were placed over the superior and inferior aspect of *tibialis anterior* using a bi-adhesive foam layer and conductive paste (Ten20, Weaver and Company, CO, USA). A dampened strap electrode (WS2, OTBioelettronica, Torino, Italy) was positioned around the ankle joint as a ground electrode. Surface electromyograms (sEMG) were recorded during the submaximal ramp-shaped contractions. The sEMG signals were acquired in monopolar mode, amplified (256×), band-pass filtered (10–500 Hz), and converted to a digital signal at 2048 Hz by a 16-bit wireless amplifier (Sessantaquattro, OTBio-elettronica, Torino, Italy) using OTBioLab+ software (version 1.3.0., OTBioelettronica, Torino, Italy) before being stored for offline analysis.

### Motor unit analyses

#### Motor unit identification

The recorded data were processed offline using the DEMUSE software (Holobar & Zazula, 2007). For each muscle from Experiment 1, only the pair of ramp-shaped contractions yielding the lowest deviation from torque trajectory were analysed for the handgrip or resting control conditions. The same was adopted on Experiment 2, in which only the motor units from one pair of ramp-shaped contractions (vibration and control conditions) yielding the lowest deviation from torque trajectory were analysed. If pairs of contractions presented a similar torque trajectory, the pair with the highest number of identified motor units was analysed. High-density sEMG signals were band-pass filtered (20–500 Hz) with a second-order, zero-lag Butterworth filter. Thereafter, a blind source separation method, the convolutive kernel compensation (CKC) method, was used for signal decomposition (Holobar & Zazula, 2007; Holobar *et al*., 2014) from each triangular contraction. CKC yields the filters of individual motor units that, when applied to high-density sEMG signals, estimate the motor unit spike trains (Holobar & Zazula, 2007; Holobar *et al*., 2014). The motor unit filters were used to identify the same motor unit across different time points. In Experiment 1, motor units were tracked before and after the handgrip or resting control conditions, but not between each condition. In Experiment 2, motor units were tracked between the vibrator on and vibrator off conditions. The motor unit filters identified by convolutive kernel compensation at individual contractions on each time point were applied to the concatenated high-density sEMG signals recorded at other time points. Afterwards, motor unit filters identified from each time point were applied to the concatenated recordings (Frančič & Holobar, 2021, 2022) yielding the motor unit spike trains of all the identified motor units across all the concatenated time points. After removing motor unit duplicates simultaneously identified from two time points a trained investigator manually inspected motor unit spike trains and edited the discharge patterns of the motor units. Only motor units visually inspected and with a pulse-to-noise ratio equal to or greater than 30 dB were kept for further analysis (Holobar *et al*., 2014).

#### Estimation of PIC strength (ΔF) and peak discharge rates

The observed discharge events for each motor unit were converted into instantaneous discharge rates and fitted into a 5^th^-order polynomial function. PIC strengths were estimated using the paired motor unit analysis (Gorassini *et al*., 2002). Motor units with a low recruitment threshold (i.e., control units) were paired with higher recruitment threshold motor units (i.e., test units). ΔF was calculated as the change in discharge rates of the control motor unit from the moment of recruitment to the moment of de-recruitment of the test unit (Gorassini *et al*., 2002). Motor unit pairs were composed of motor units with a rate-to-rate correlations between the smoothed discharge rate polynomials of r ≥ 0.7 and the test units were recruited at least 1.0 s after the control units (Gorassini *et al*., 2002; Hassan *et al*., 2020). ΔFs obtained for each control unit were averaged to obtain a single ΔF for each test motor unit. ΔFs for the motor units tracked over time were derived from the same pairs of motor units on each condition (i.e., Experiment 1, between before and after handgrip or resting control conditions, and Experiment 2, between vibrator on and vibrator off conditions). The maximum value obtained from the polynomial curve was considered the peak discharge rate. The relative torque (%) produced at the time in each motor unit was recruited was considered the recruitment threshold. Figure 2 shows an example of paired motor unit analysis used to calculate ΔF from a single test unit using three control units.

**Figure 2.**
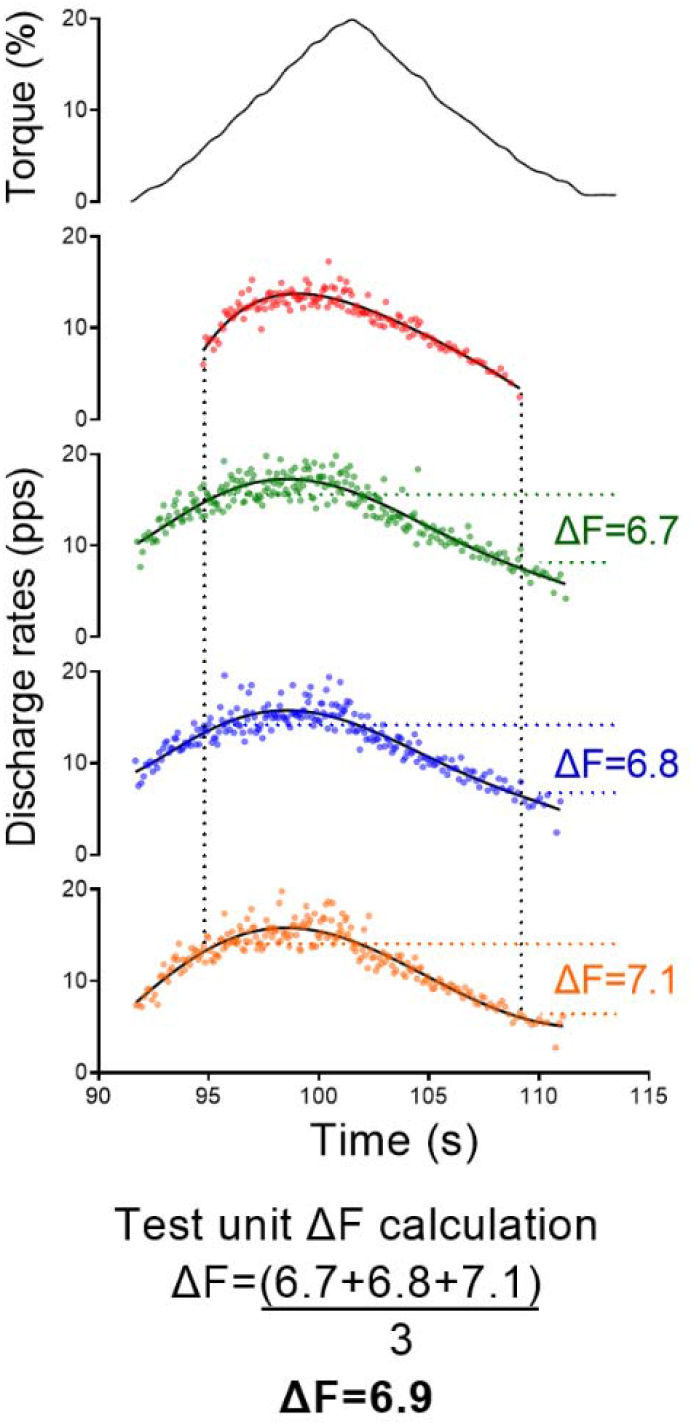
Data illustrating the ΔF calculation using paired motor unit analysis. The top panel shows the torque traces for contractions to 20% of the participant’s dorsiflexion maximal voluntary torque. The subsequent panels display a *tibialis anterior* test motor unit (red colour) and three control units (green, blue, and orange colours). The black continuous lines are the 5th-order polynomial fits for the control units. The ΔFs obtained from each control unit were averaged, resulting in a single value for the test unit.

**Figure 3.**
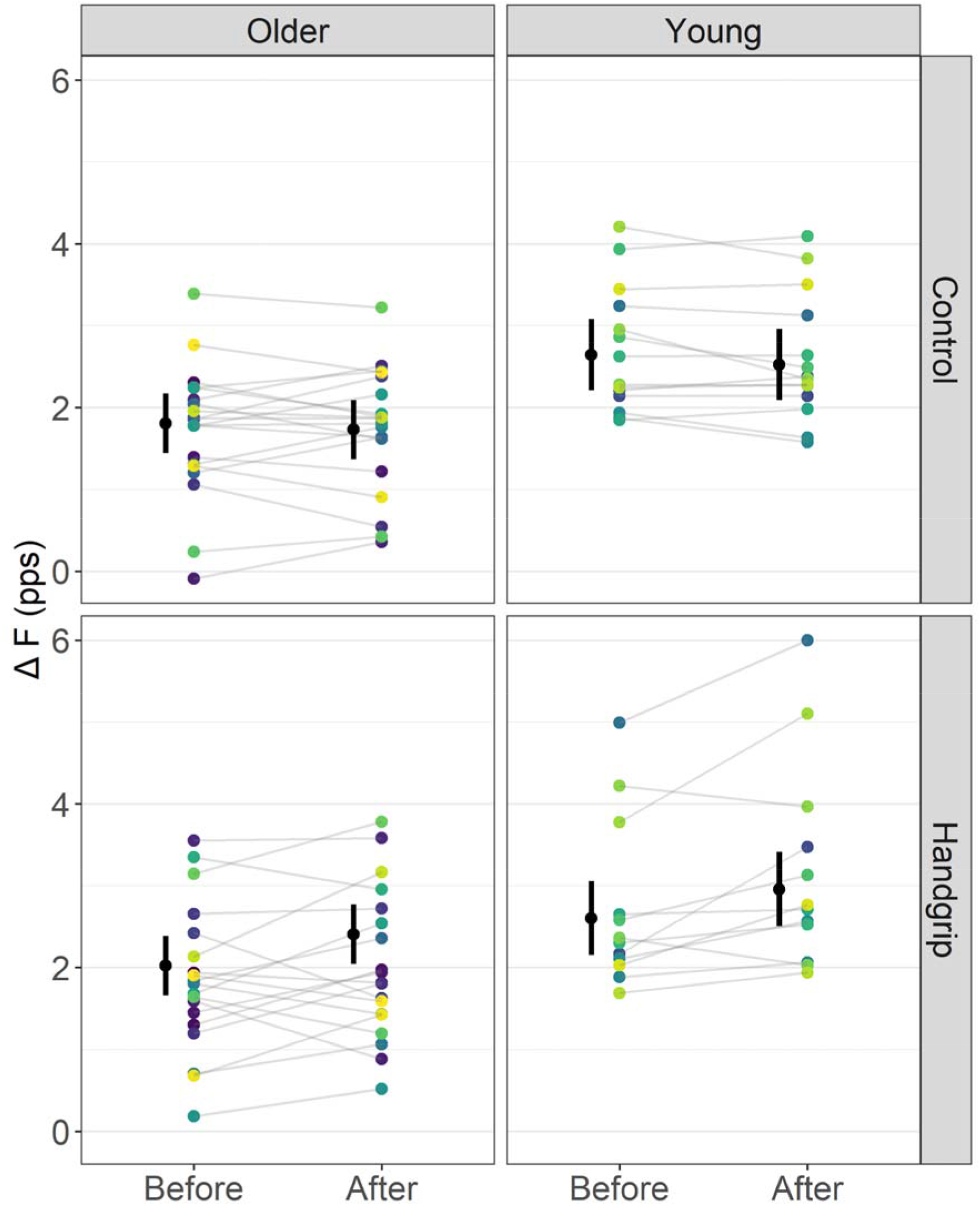
*Soleus* ΔF outcomes in control and handgrip conditions for older and young adults. Note that ΔF remained unchanged before and after the control condition but increased in the handgrip condition in both older and young adults. The mean (black circle) and 95% confidence intervals are offset to the left. Individual data points (average ΔF per participant) are coloured by participants. pps = peaks per second.

**Figure 4.**
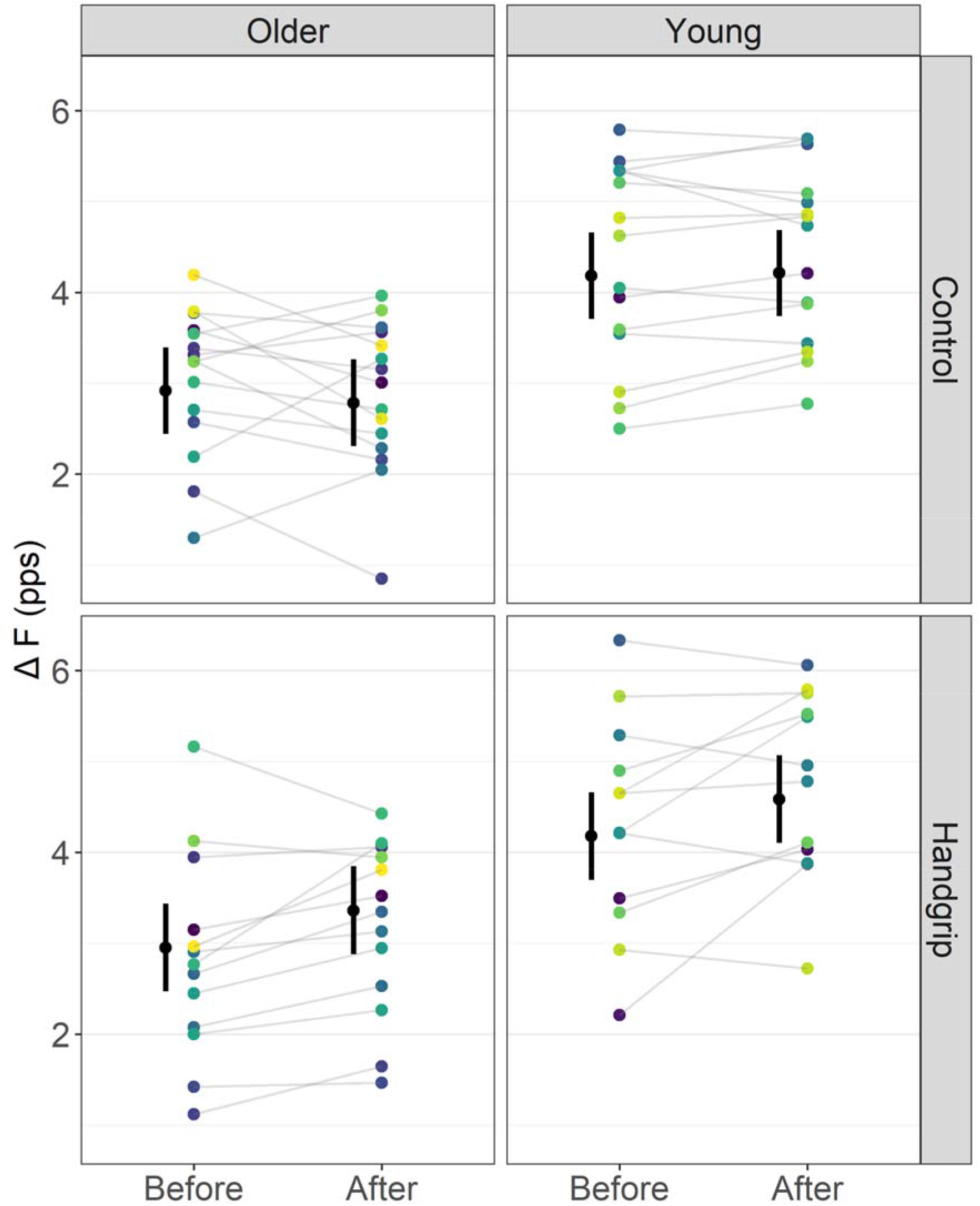
*Tibialis anterior* ΔF outcomes in control and handgrip conditions for older and young adults. Note that ΔF remained unchanged before and after the control condition but increased in the handgrip condition in both older and young adults. The mean (black circle) and 95% confidence intervals are offset to the left. Individual data points (average ΔF per participant) are coloured by participants. pps = peaks per second.

**Figure 5.**
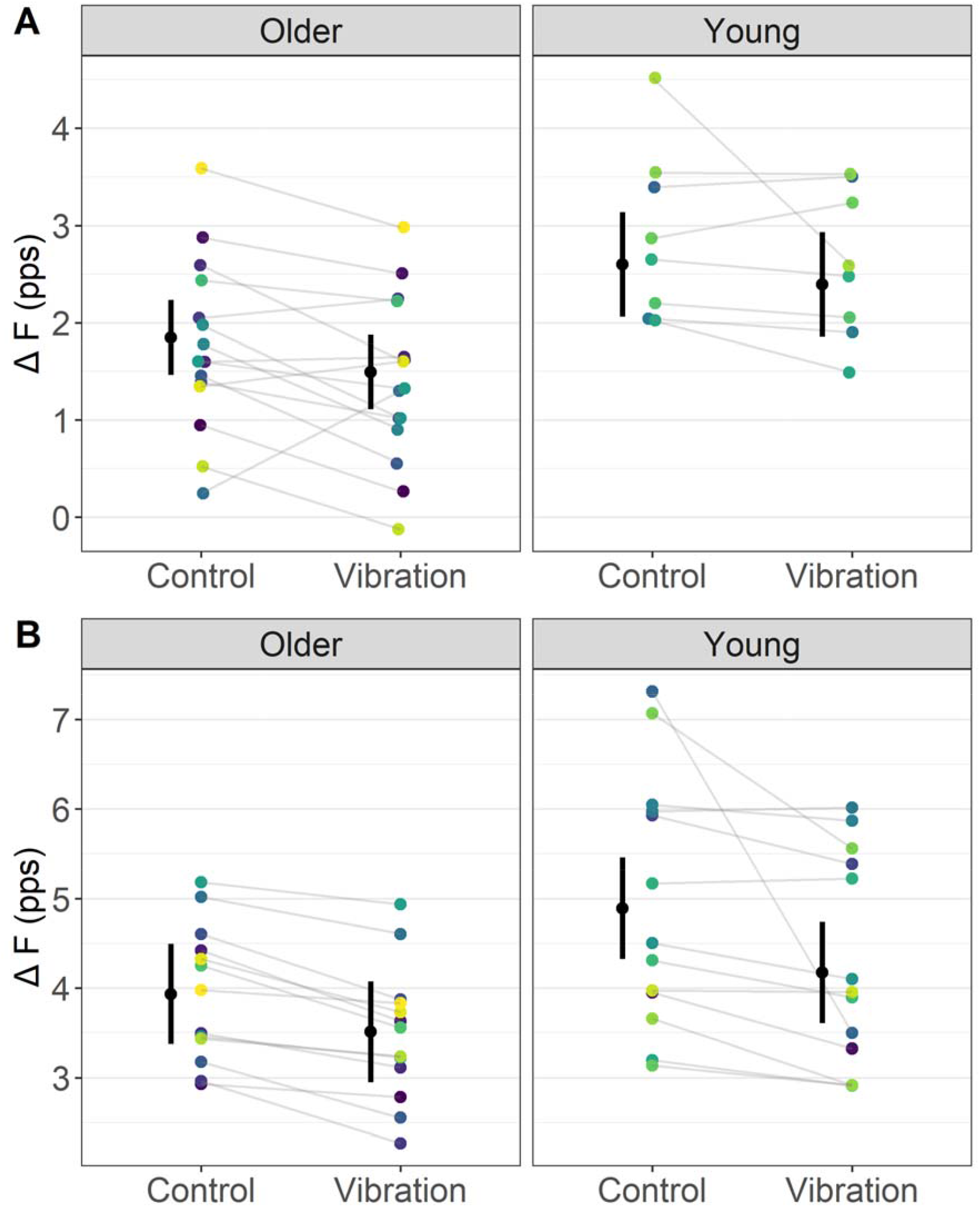
*Soleus* (Panel A) and *tibialis anterior* (Panel B) ΔF outcomes in control and vibration conditions for older and young adults. Note that, for *soleus*, ΔF was reduced during the vibration compared to the control condition, similarly between older and young adults. For *tibialis anterior*, ΔF was reduced in older adults during the vibration condition, but to a smaller magnitude than young adults (i.e., condition by age interaction). The mean (black circle) and 95% confidence interval are offset to the left. Individual data points (average ΔF per participant) are coloured by participants. pps = peaks per second.

### Data analyses

In Experiment 1, separate linear mixed-effect models were used for each muscle to compare ΔF, peak discharge rates, and recruitment thresholds between conditions (i.e., control and handgrip), age groups (i.e., older and young adults), and over time (i.e., before and after handgrip or control conditions), used as a fixed factors (Yu *et al*., 2021). Additional linear mixed-effect models were used to compare ΔF mean differences (after – before handgrip or control conditions) between conditions and age groups to remove the time factor from the model, and results are presented in Supplementary Material 1. In Experiment 2, linear mixed-effect models were used to compare ΔF, peak discharge rates, and recruitment thresholds between conditions (i.e., control and vibrator), used as a fixed factors (Yu *et al*., 2021). For both experiments, *soleus* and *tibialis anterior* were fitted to separate statistical models. Single motor units were treated as repeated measures, nested according to each participant, and a random intercept was included for each participant in the study to account for the correlation between repeated observations on each individual (i.e., 1| participant/motor unit ID). When a significant effect was observed, Bonferroni post-hoc correction was adopted for pairwise comparison. The effect sizes derived from the F ratios were calculated with the omega squared (ω^2^) method (0–0.01, very small; 0.01–0.06, small; 0.06–0.14, moderate; and >0.14, large) (Lakens, 2013). Intraclass correlation coefficient (ICC) and standard error of measurement (SEM) were calculated between ΔF before and after the control condition from Experiment 1. The standardised difference (Cohen’s *d*) between time points was also calculated using the population standard deviation from each respective linear mixed-effects model as the denominator (Lenth *et al*., 2021). All analyses were completed in R (version 4.0.5) using the RStudio environment (version 1.4.1717). Linear mixed-effects models were fitted using the lmerTest package (Kuznetsova *et al*., 2017). Estimated marginal mean differences and 95% confidence intervals between time points were determined using the emmeans package (Lenth *et al*., 2021). Significant difference was accepted at p ≤ 0.05. All descriptive data are presented as mean (95% confidence interval lower and upper limits), unless indicated differently. The dataset and R code can be found at https://github.com/orssatto/PICs-ageing_2.0.

## RESULTS

### Experiment 1

In summary, ΔF and peak discharge rates were increased in motor units of young and older adults following the handgrip condition but not the control condition. These results were observed in both *soleus* and *tibialis anterior*, independent of age. Recruitment threshold did not change in *soleus*, independent of age, but increased in *tibialis anterior* in older adults only. Descriptive statistics for each group, condition, time point, and muscle are presented in Table 1.

**Table 1.** Estimated marginal means and mean differences (95% confidence interval lower and upper limits) for ΔF, peak discharge rates, and recruitment thresholds before and after control or handgrip conditions in young and older adults and for *soleus* and *tibialis anterior*.

| Variables | $\Delta F$ (pps) | Peak discharge<br>rates (pps) | Recruitment<br>threshold (% of peak<br>torque) |
| --- | --- | --- | --- |
| **Older adults - *Soleus*** |  |  |  |
| Before control | 1.81 (1.44, 2.17) | 8.75 (8.17, 9.32) | 8.11 (6.92, 9.30) |
| After control | 1.73 (1.36, 2.09) | 8.62 (8.05, 9.20) | 8.39 (7.20, 9.57) |
| After – before control | -0.08 (-0.37, 0.21) | -0.13 (-0.39, 0.14) | 0.28 (-0.76, 1.31) |
| Before handgrip | 2.02 (1.66, 2.39) | 8.54 (7.96, 9.12) | 8.66 (7.47, 9.85) |
| After handgrip | 2.41 (2.04, 2.77) | 9.00 (8.42, 9.57) | 9.01 (7.82, 10.20) |
| After – before handgrip | 0.39 (0.10, 0.67) | 0.46 (0.20, 0.72) | 0.35 (-0.68, 1.38) |
| **Young adults - *Soleus*** |  |  |  |
| Before control | 2.64 (2.21, 3.08) | 10.65 (9.95, 11.36) | 9.20 (7.79, 10.61) |
| After control | 2.53 (2.09, 2.96) | 10.75 (10.04, 11.45) | 9.03 (7.62, 10.45) |
| After – before control | -0.12 (-0.44, 0.20) | 0.10 (-0.19, 0.39) | -0.17 (-1.31, 0.98) |
| Before handgrip | 2.60 (2.15, 3.06) | 10.57 (9.85, 11.28) | 9.54 (8.06, 11.02) |
| After handgrip | 2.96 (2.51, 3.41) | 10.91 (10.20, 11.63) | 8.42 (6.94, 9.90) |
| After – before handgrip | 0.35 (-0.04, 0.74) | 0.34 (-0.01, 0.70) | -1.12 (-2.51, 0.28) |
| **Older adults – *Tibialis anterior*** |  |  |  |
| Before control | 2.92 (2.44, 3.40) | 12.14 (11.09, 13.19) | 10.39 (8.94, 11.84) |
| After control | 2.79 (2.31, 3.26) | 11.89 (10.84, 12.95) | 10.88 (9.43, 12.33) |
| After – before control | -0.13 (-0.39, 0.13) | -0.25 (-0.60, 0.10) | 0.49 (-0.17, 1.15) |
| Before handgrip | 2.95 (2.47, 3.44) | 12.04 (10.98, 13.10) | 9.71 (8.24, 11.18) |
| After handgrip | 3.36 (2.88, 3.85) | 12.22 (11.16, 13.28) | 11.36 (9.88, 12.83) |
| After - before handgrip | 0.41 (0.10, 0.72) | 0.18 (-0.23, 0.59) | 1.65 (0.80, 2.49) |
| **Young adults - *Tibialis anterior*** |  |  |  |
| Before control | 4.18 (3.71, 4.66) | 14.75 (13.71, 15.79) | 11.36 (9.91, 12.80) |
| After control | 4.21 (3.74, 4.69) | 14.71 (13.67, 15.75) | 10.79 (9.34, 12.23) |
| After – before control | 0.03 (-0.25, 0.30) | -0.04 (-0.41, 0.33) | -0.57 (-1.29, 0.15) |
| Before handgrip | 4.18 (3.70, 4.66) | 14.12 (13.08, 15.17) | 11.56 (10.10, 13.02) |
| After handgrip | 4.59 (4.10, 5.07) | 14.66 (13.62, 15.75) | 11.05 (9.59, 12.51) |
| After – before handgrip | 0.41 (0.10, 0.71) | 0.53 (0.12, 0.94) | -0.51 (-1.31, 0.29) |

#### Estimates of PICs (ΔF)

For *soleus*, a time by condition interaction on ΔF (F = 19.27, ω^2^ = 0.05, p < 0.001), but not a time by condition by age interaction (F < 0.01, ω^2^ < 0.001, p = 0.967), was detected. ΔF did not change over time in the control condition [mean difference (md) = -0.10 (−0.28, 0.09) pps, *d* = -0.17 (−0.41, 0.08), p = 0.17] but increased in the handgrip condition [md = 0.37 (0.16, 0.57) pps, *d* = 0.62 (0.35, 0.90), p < 0.001]. Before the interventions, ΔF was not different between the control and handgrip conditions [md = -0.09 (−0.30, 0.13), *d* = -0.15 (−0.43, 0.14), p = 0.292]. However, ΔF was higher in the handgrip condition than the control condition after the intervention [md = 0.55 (0.34, 0.77), *d* = 0.932 (0.64, 1.22), p < 0.001]. ICC for ΔF measured before and after the control condition was 0.939 (0.893, 0.965) and SEM was 0.219 pps.

For *tibialis anterior*, there was a time by condition interaction on ΔF for *tibialis anterior* (F = 23.81, ω^2^ = 0.03, p < 0.001) but not a time by condition by age interaction (F = 0.76, ω^2^ < 0.001, p = 0.385). ΔF did not change over time in the control condition [md = -0.05 (−0.21, 0.11) pps, d = -0.07 (−0.24, 0.10), p = 0.399], but increased in the handgrip condition [md = 0.42 (0.23, 0.59) pps, *d* = 0.55 (0.35, 0.74), p < 0.001]. Before the interventions, ΔF was not different between control and handgrip conditions [md = -0.01 (−0.20, 0.17) pps, *d* = -0.02 (−0.22, 0.18), p = 0.838]. However, ΔF was higher in the handgrip than the control condition after the interventions [md = 0.64 (0.44, 0.84) pps, *d* = 0.61 (0.40, 0.81), p < 0.001]. ICC for ΔF measured before and ΔF after the control condition was 0.898 (0.819, 0.944) and SEM was 0.372 pps.

#### Peak discharge rates

For *soleus*, a time by condition interaction (F = 18.52, ω^2^ = 0.05, p < 0.001), but not a time by condition by age interaction (F = 3.02, ω^2^ < 0.001, p = 0.083), was detected. Peak discharge rates did not change over time in the control condition [md = -0.02 (−0.18, 0.15) pps, *d* = - 0.03 (−0.27, 0.21), p = 0.812] but increased in the handgrip condition [md = 0.40 (0.21, 0.59) pps, *d* = 0.74 (0.47, 1.02), p < 0.001]. For *tibialis anterior*, there was a time by condition interaction (F = 15.57, ω^2^ = 0.02, p < 0.001) but no time by condition by age interaction (F = 0.345, ω^2^ < 0.001, p = 0.557). Peak discharge rates did not change over time in the control condition (md = -0.14 (−0.36, 0.07) pps, d = -0.14 (−0.31, 0.03), p = 0.084) but increased after the handgrip condition (md = 0.36 (0.11, 0.60) pps, d = 0.36 (0.16, 0.56), p = 0.002), irrespective of age.

#### Recruitment thresholds

For *soleus*, there was no time by condition by age interaction (F = 1.81, ω^2^ < 0.001, p = 0.180) nor time by condition interaction (F = 1.33, ω^2^ < 0.001, p = 0.250). Recruitment threshold remained unchanged before and in both the control (md = 0.06 (−0.60, 0.71) % of peak torque, *d* = 0.03 (−0.28, 0.28), p = 0.826) and handgrip (md = -0.44 (−1.2, 0.32) % of peak torque, *d* = -0.18 (−0.45, 0.09), p = 0.179) conditions, irrespective of age. For *tibialis anterior*, there was a time by condition by age interaction (F = 4.79, ω^2^ < 0.001, p = 0.029). Recruitment threshold increased after handgrip in older [*d* = 0.84 (0.55, 1.13), p < 0.001] but not for young adults [*d* = -0.26 (−0.53, 0.01), p = 0.527], and remained unchanged before and after control for both older (p = 0.319) and young adults (p = 0.235).

### Experiment 2

In summary, ΔF decreased in motor units of both young and older adults when vibration was applied to the antagonist tendon. In *soleus*, these reductions were similar for young and older adults but in *tibialis anterior* the magnitude of reduction was smaller for older than young adults. Peak discharge rates remained unchanged in *soleus* but reduced for *tibialis anterior*, irrespective of age. Recruitment threshold remained unchanged in *soleus*, irrespective of age, but increased for young adults only in *tibialis anterior*. Descriptive statistics for each group, condition, and muscle are presented in Table 2.

**Table 2.**
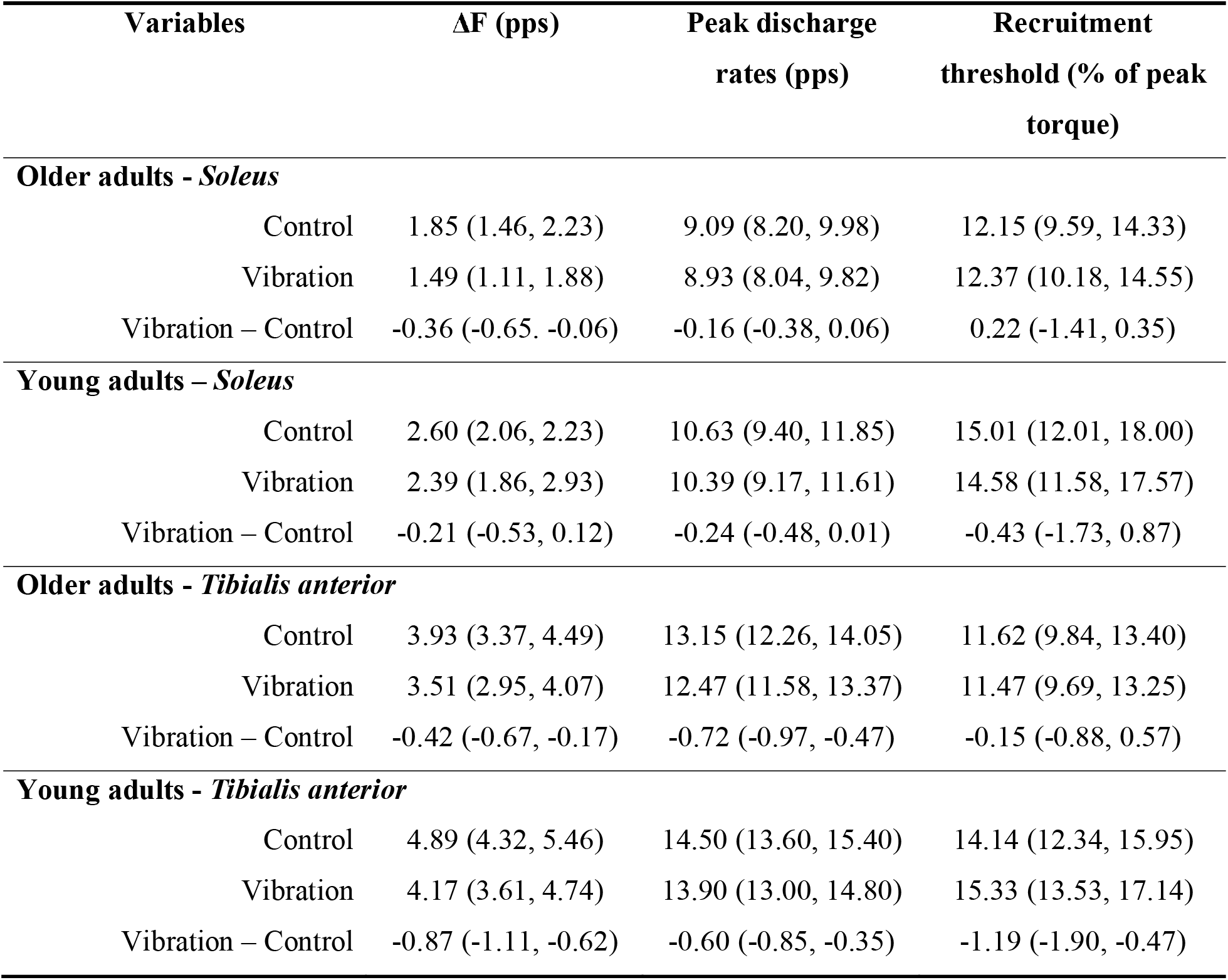
Estimated marginal means and mean differences (95% confidence interval lower and upper limits) for ΔF, peak discharge rates, and recruitment thresholds for vibration and control conditions.

| Variables | $\Delta F$ (pps) | Peak discharge rates (pps) | Recruitment threshold (% of peak torque) |
| --- | --- | --- | --- |
| **Older adults - *Soleus*** |  |  |  |
| Control | 1.85 (1.46, 2.23) | 9.09 (8.20, 9.98) | 12.15 (9.59, 14.33) |
| Vibration | 1.49 (1.11, 1.88) | 8.93 (8.04, 9.82) | 12.37 (10.18, 14.55) |
| Vibration – Control | -0.36 (-0.65, -0.06) | -0.16 (-0.38, 0.06) | 0.22 (-1.41, 0.35) |
| **Young adults – *Soleus*** |  |  |  |
| Control | 2.60 (2.06, 2.23) | 10.63 (9.40, 11.85) | 15.01 (12.01, 18.00) |
| Vibration | 2.39 (1.86, 2.93) | 10.39 (9.17, 11.61) | 14.58 (11.58, 17.57) |
| Vibration – Control | -0.21 (-0.53, 0.12) | -0.24 (-0.48, 0.01) | -0.43 (-1.73, 0.87) |
| **Older adults - *Tibialis anterior*** |  |  |  |
| Control | 3.93 (3.37, 4.49) | 13.15 (12.26, 14.05) | 11.62 (9.84, 13.40) |
| Vibration | 3.51 (2.95, 4.07) | 12.47 (11.58, 13.37) | 11.47 (9.69, 13.25) |
| Vibration – Control | -0.42 (-0.67, -0.17) | -0.72 (-0.97, -0.47) | -0.15 (-0.88, 0.57) |
| **Young adults - *Tibialis anterior*** |  |  |  |
| Control | 4.89 (4.32, 5.46) | 14.50 (13.60, 15.40) | 14.14 (12.34, 15.95) |
| Vibration | 4.17 (3.61, 4.74) | 13.90 (13.00, 14.80) | 15.33 (13.53, 17.14) |
| Vibration – Control | -0.87 (-1.11, -0.62) | -0.60 (-0.85, -0.35) | -1.19 (-1.90, -0.47) |

#### Estimates of PICs (ΔF)

For *soleus*, a condition effect on ΔF (F = 11.16, ω^2^ = 0.12, p = 0.001), but not a condition by age interaction (F = 0.80, ω^2^ < 0.001, p = 0.374), was detected. ΔF was lower in the vibration than the control condition, irrespective of age group [md = 0.28 (0.11, 0.45) pps, *d* = 0.54 (0.19, 0.89)]. For *tibialis anterior*, there was a condition by age interaction on ΔF (F = 4.66, ω^2^ = 0.01, p = 0.032). ΔF reductions in older adults [md = 0.42 (0.17, 0.67) pps, *d* = 0.51 (0.26, 0.76)] were smaller than in young adults [md = 0.72 (0.47, 0.97) pps, *d* = 0.87 (0.62, 1.11)]. A pair of outlier motor units was identified in the *tibialis anterior* analysis but excluding them from the model did not remove the evidence of condition by age interaction (F = 3.68, ω^2^ = 0.01, p = 0.056).

#### Peak discharge rates and recruitment threshold

For *soleus*, a condition effect on peak discharge rates (F = 9.98, ω^2^ = 0.10, p = 0.002), but not a condition by age interaction (F = 0.41, ω^2^ < 0.001, p = 0.526), was detected. Peak discharge rates were 0.20 (0.07, 0.32) pps lower in the vibration than the control condition [*d* = 0.51 (0.17, 0.86)], irrespective of age. There was no condition effect (F = 0.10, ω^2^ = 0.01, p = 0.752) nor a condition by age interaction on recruitment threshold (F = 0.95, ω^2^ < 0.001, p = 0.333). For *tibialis anterior*, a condition effect on peak discharge rates (F = 86.22, ω^2^ = 0.23, p < 0.001), but not a condition by age interaction (F = 0.34, ω^2^ < 0.001, p = 0.559), was detected. Peak discharge rates were 0.64 (0.50, 0.78) pps lower in the vibration than control condition (*d* = 0.77 (0.59, 0.95)) irrespective of age. There was a condition by age interaction on recruitment threshold (F = 11.61, ω^2^ = 0.04, p < 0.001). Recruitment threshold remained unchanged in older adults (*d* = -0.06 (−0.31, 0.18), p = 0.585) but increased in young adults (*d* = 0.50 (0.26, 0.74), p < 0.001).

### Motor unit identification

In Experiment 1, 201 *soleus* motor units for 21 older adults and 110 *soleus* motor units for 14 older adults were identified by the decomposition algorithm that could be matched before and after the control condition, resulting in 78 and 64 test units, respectively. 187 *soleus* motor units were matched before and after the handgrip contraction for 19 older adults and 101 for 12 young adults, resulting in 79 and 43 test units, respectively. 326 *tibialis anterior* motor units for 16 older adults and 262 for 15 young adults were matched before and after the control condition, resulting in 154 and 132 test units, respectively. 253 *tibialis anterior* motor units for 14 older adults and 222 for 13 young adults were matched before and after the control condition, resulting in 110 and 116 test units, respectively. In Experiment 2, 97 *soleus* motor units for 15 older adults and 60 motor units for 8 young adults were matched between control and vibration conditions, resulting in 42 and 35 test units respectively. For, 246 *tibialis anterior* motor units for 13 older adults and 215 motor units from 13 young adults were matched between control and vibration conditions, resulting in 143 and 147 test units, respectively. Motor units sample median and interquartile range per experiment, group, and conditions are presented in Supplementary material 2.

## DISCUSSION

This study tested the ability of older and young adults to both amplify and inhibit motoneuronal PICs, estimated using paired motor unit analyses (i.e., ΔF), in response to a remote handgrip contraction and to reciprocal inhibition, respectively. We previously showed that ΔF was lower in older than young adults, measured in the same cohort of subjects (Orssatto *et al*., 2021*a*)). However, here we present the first evidence that the abilities of older adults to modulate PICs in response to a remote handgrip contraction, which theoretically increases monoaminergic projections onto the motor neurones, is preserved in both *soleus* and *tibialis anterior*. These results are not consistent with our hypothesis that older adults would have a reduced ability to increase ΔF after a remote handgrip contraction. However, we also observed that older adults’ abilities to inhibit ΔF was preserved in *soleus* but not *tibialis anterior*, in which they presented attenuated inhibition. These data partially agree with our initial hypothesis that inhibition would be impaired in older adults but clarify that the effect may not be consistent between muscles or muscle groups, which we discuss further below. The novel findings from this study are fundamental to our understanding of facilitatory-inhibitory control of PICs in older and young adults.

### Experiment 1 – Estimates of PIC neuromodulation

Increases in ΔF following the 30-s remote handgrip contraction (at 40% of maximal force) for both *soleus* (16%) and *tibialis anterior* (11.4%) were consistent with our initial hypothesis. This amplification likely results from the increased available serotonin delivered by descending tracts within the spinal cord, which has diffuse input onto motor neurones across diverse muscle groups (Heckman *et al*., 2008; Wei *et al*., 2014). This should be predominately attributed to the activity of serotonergic projections to the spinal cord, which is increased along with motor output (Jacobs *et al*., 2002), than noradrenaline, which is more influenced by arousal state (Aston-Jones *et al*., 2001). Therefore, the increased availability of serotonin along with its highly diffuse descending projections into the spinal cord is expected to be the main mechanism underpinning the observed PIC amplification.

Contrary to expectation, young and older adults showed similar PIC amplification after the sustained handgrip contraction in both *soleus* and *tibialis anterior*. One consideration is that this similarity might be limited to relatively light submaximal contractions, such as the ramp contractions in our study which peaked at 20% of maximum torque capacity. The choice of this torque level was largely based on matching the forces that may be exerted during walking or postural sway, and partly based on ensuring optimal motor unit identification (fewer motor unit pairs are detected at higher contraction levels and fatigue is generated by prolonged and high intensity contractions, making the technique less feasible as contraction level increases). However, the magnitude of PIC amplification depends critically on the level of monoaminergic drive onto the motor neurones (Hounsgaard *et al*., 1988; Lee & Heckman, 1999, 2000), which in turn is affected by the exerted contraction intensity (Jacobs *et al*., 2002). Thus, stronger contractions demand greater release of monoamines onto the motor neurones and should enable greater PIC amplification (Hounsgaard *et al*., 1988; Lee & Heckman, 1999, 2000; Orssatto *et al*., 2021*b*). Using contractions to 20% of maximum will have prompted only a moderate increase in serotonin concentration, and potentially below some existing ceiling, allowing the observed PIC amplification in the older group to occur subsequent to the handgrip contraction. We cannot rule out the possibility that higher intensity contractions, providing stronger monoaminergic input to amplify PICs, would have reduced the capacity for the handgrip contraction itself to provide monoaminergic drive above that already provided by the contraction itself, in either the older or young adults. Nonetheless, a recent study by our group (including a subset of older adults from the present study) showed that non-strength-trained older adults were unable to further amplify *soleus* PICs as plantar flexion contraction intensity increased from 20 to 40% of peak torque (Orssatto *et al*., 2022*b*). This is not consistent with the behaviour of increased PICs amplification observed in young adults when contraction intensity increased from 10 to 30% of peak torque (Orssatto *et al*., 2021*b*), indicating an age-dependent difference in the capacity to increase PIC strength with contraction intensity. These results are suggestive of a potentially impaired serotonergic input onto the motor neurones that may manifest at higher contraction intensities in older individuals only. Although ΔF values are more difficult to attain in higher-force contractions, it would be of interest to test PIC amplification at higher force levels in future experiments.

### Experiment 2 – Estimates of PIC inhibition

We tested the hypothesis that ΔF should decrease during reciprocal inhibition, induced by antagonist tendon vibration, in *soleus* and *tibialis anterior*. Our data demonstrate that high-frequency vibration of *tibialis anterior* and the Achilles tendon successfully induced reciprocal inhibition, reducing ΔF in *soleus* by 0.28 pps (12.6%) in both older and young adults, and in *tibialis anterior* by 0.42 pps (10.7%) in older adults and 0.87 pps (17.8%) in young adults. High-frequency tendon vibration triggers excitatory drive from Ia afferents via stimulation of muscle spindles, which consequently generates reciprocal inhibition by activating the Ia inhibitory inter-neurones (Burke *et al*., 1976). The reductions in ΔF during the reciprocal inhibition condition shown in our data are consistent with findings from animal experiments showing reciprocal inhibition-evoked PIC reductions induced by *in-vivo* voltage-clamp techniques in cats (Kuo *et al*., 2003; Hyngstrom *et al*., 2007). However, using such techniques, dendritic PICs were reduced by ∼50% in the ankle extensors by imposition of small joint rotations (Hyngstrom *et al*., 2007) and by ∼69% during tonic nerve electrical stimulation to antagonist muscles (in a standard monoaminergic state) (Kuo *et al*., 2003). Both of these animal experiments therefore showed a much greater reduction than we observed in the current experiment. Nonetheless, our results are consistent with recent human experiments showing comparable ΔF reductions of 0.54 ± 0.09 pps in both *tibialis anterior* (effect size g = 0.49) and *soleus* (g = 0.26) using a similar protocol of antagonist tendon vibration (Pearcey *et al*., 2022). We also recently observed similar ΔF reductions of 0.33 pps (9.8%) in *gastrocnemius medialis* of young adults after reciprocal inhibition was induced by electrical stimulation of the common peroneal nerve (Mesquita *et al*., 2022). In addition, decreasing the level of reciprocal inhibition (applying stimulations to the common peroneal nerve, below motor unit threshold) increased ΔF in *soleus* (Vandenberk & Kalmar, 2014), and artificially activating an inhibitory reflex by sural nerve stimulation reduced the initial steep increases in discharge rates in *tibialis anterior* motor neurones during ramped contractions, which is likely modulated by PICs (Revill & Fuglevand, 2017). These data suggest that the magnitude of reciprocal inhibition on PICs may be smaller when assessed by the ΔF method in humans than in direct measurements in animal experiments. This could result from the stronger reciprocal inhibition evoked in animal experiments (Kuo *et al*., 2003; Hyngstrom *et al*., 2007) and/or because ΔF only estimates the portion of PICs above the discharging threshold (Gorassini *et al*., 2002).

We hypothesised that older adults would present smaller ΔF reductions than young adults because reciprocal inhibition is generally considered to be reduced with ageing in both *soleus* and *tibialis anterior* (Kido *et al*., 2004). However, our data show a difference only in *tibialis anterior*. The weaker inhibition of *tibialis anterior* in the older adults might be explained by (i) an age-related alteration in reciprocal inhibition due to a decrease in muscle spindle quantity or changes in their structure, innervation, and sensitivity (Henry & Baudry, 2019), (ii) by reductions in the total number of nerve fibres, including spindle afferents (Swallow, 1966), and deterioration of Ia afferent pathways (observed in aged rats) (Vaughan *et al*., 2017), (iii) by changes in the transmission efficacy at the Ia inhibitory inter neurone, or (iv) by efferent fibre impairment (Geertsen *et al*., 2017). Although there is a lack of consensus in the literature, reduced transmission efficacy of Ia afferents with ageing (measured indirectly with h-reflex responses) (Scaglioni *et al*., 2003, 2012; Klass *et al*., 2011; Škarabot *et al*., 2019, 2020) could contribute to an impaired reciprocal inhibition of motor neuronal PICs. As a consequence, this could partly underpin the increased *tibialis anterior* coactivation during important daily tasks such as single leg stance (Kurz *et al*., 2018) and during gait (Hortobágyi *et al*., 2009) in older adults, which should be explored in future studies.

The dissimilar response between *soleus* and *tibialis anterior* may speculatively be explained by distinct, between-muscle effects of ageing on muscle spindles and sensory afferents. Although no direct comparison between *soleus* and *tibialis anterior* exists, reductions in muscle spindle diameter have been observed in aged *deltoid* and *extensor digitorum*, although not in *quadriceps* or *biceps brachii*, and decreases in the number of intrafusal fibres in *deltoid* have also been detected (Kararizou *et al*., 2005). It is therefore possible that ageing differently affects *soleus* and *tibialis anterior* muscle spindles and sensory afferents, and this might be confirmed in future experiments. Interestingly, passive ankle plantar and dorsiflexion did detectibly influenced cortico-spinal responses in *tibialis anterior* in young adults but not in older adults, while it remained unchanged in *soleus* in both groups (Škarabot *et al*., 2020). These data support the assertion that *soleus* and *tibialis anterior* afferent and/or efferent pathways might be differently affected by ageing. Also, agerelated decreases in vibration–electromyogram coherence, response gain, response amplitude, and scaling of the *soleus* response to Achilles tendon stimuli have been shown (Mildren *et al*., 2020), indicating impaired proprioceptive feedback from *soleus*. The different functional roles of *soleus* and *tibialis anterior* in daily tasks should also be considered when interpreting the present data. Both muscles are active during daily living activities, such as upright standing and locomotor propulsion (e.g., gait) (Soames & Atha, 1981; Masani *et al*., 2013), however *soleus* serves an anti-gravity role, implying that motor units are active for longer and produce greater cumulative force than flexor muscles during daily living activities. Additional data supporting this claim includes evidence that peak discharging rates are maintained with ageing in well-used muscles (e.g., hand muscles, *quadriceps*, and *triceps surae*) but decline markedly in lesser used muscles (e.g., *hamstrings* and *tibialis anterior*) (Orssatto *et al*., 2022*a*). In fact, studies show that disuse can aggravate the deleterious effects of ageing on the nervous system, while trained older adults show a substantial preservation of neural function (Aagaard *et al*., 2010; Mcgregor *et al*., 2011; Unhjem *et al*., 2016; Hvid *et al*., 2018; Orssatto *et al*., 2020). Regardless, direct comparisons between *soleus* and *tibialis anterior* are required to elucidate the mechanisms underpinning the dissimilar effects of ageing on PIC inhibitory control between *soleus* and *tibialis anterior*.

### Strengths, limitations and delimitations

Our study employed a validated and extensively used method of PIC strength estimation in humans (Udina *et al*., 2010; Hassan *et al*., 2019, 2021; Trajano *et al*., 2020; Orssatto *et al*., 2021*b*, 2021*a*; Mesquita *et al*., 2022) that shows very good repeated-measures reliability, as demonstrated by the high reliability of measurements in the control condition of Experiment 1. However, our findings should not be extrapolated to contraction intensities and motor units with recruitment threshold higher than those used in our study (20% and 30% of peak torques), as discussed previously. Also, our findings relate only to *soleus* and *tibialis anterior* as, for example, different muscles have distinct muscle spindle concentrations (Banks, 2006), which may alter the strength of reciprocal inhibition and the effects of ageing. Daily activities require the activation of different muscle groups; thus, future investigations of the effects of aging on neuromodulatory-inhibitory control of PICs in other muscles, and its influence on possible impairments in physical function, require additional study. Lastly, we have included only non-sarcopaenic older adults, so our results should not be extrapolated to populations with different physical characteristics or potential health conditions, such as the very old (> 85 years old), sarcopenic, or frail, or those with neurological disorders or individuals with physical activity levels.

## Conclusions

We present novel data demonstrating the control of motor neuronal PIC facilitation and inhibition in older adults by estimating PIC strengths using the paired-motor unit technique. We show that older adults have a preserved ability to amplify PICs through remote muscle contraction (e.g., handgrip, as used presently), which would diffusely increase serotonergic input onto motor neurones in both *soleus* and *tibialis anterior*. Subsequently, we present evidence of age-related reciprocal inhibition-induced impairment of PIC deactivation in *tibialis anterior*, which, conversely, was preserved with ageing in *soleus*. These findings relate to tests completed in mostly lower-threshold motor units during low intensity contractions (up to 20 or 30% of the individuals’ maximal capacity) and should not be extrapolated to higher threshold motor units or higher intensity contractions. The logical next steps are to explore (1) facilitation-inhibition control of PICs in higher intensity contractions and in different muscles, (2) the variable control of PICs on motor performance, and (3) responses to exercise interventions in older adults and other clinical populations.

## ACKNOWLEDGEMENTS

The writing period of the present manuscript was funded by the Faculty Write-up Scholarship provided by Queensland University of Technology. The authors thank Dr Raphael L. Sakugawa for developing a MATLAB script that facilitated calculation of ΔF, peak discharge rates, and recruitment thresholds.

---

Supplementary Material 1.

In *soleus*, no condition by age interaction (F = 0.01, p = 0.960) or age effect (F = 0.16, p = 0.685) on ΔF mean differences was found. However, there was a condition effect (F = 28.65, p < 0.001) in which handgrip mean differences [0.370 (0.237, 0.503) pps] were higher than in control [-0.098 (0.237, 0.503) pps]. For *tibialis anterior*, no condition by age interaction (F = 1.35, p = 0.247) or age effect (F = 0.96, p = 0.336) on ΔF mean differences was found. However, there was a condition effect (F = 29.74, p < 0.001) in which handgrip mean differences [0.404 (0.251, 0.556) pps) were higher than in control [-0.06 (−0.196, 0.081) pps]. These findings agree with the output of the 3-way linear mixed models presented within the main manuscript, confirming the lack of age effect in the condition by time interaction for both *soleus* and *tibialis anterior*.

**Supplementary Material 2.** Total and test motor unit sample medians (interquartile range) for each group and condition used in Experiments 1 and 2.

| Group | Older adults |  | Young adults |  |
| --- | --- | --- | --- | --- |
| Condition | Control | Handgrip | Control | Handgrip |
| *Soleus* motor units | 8 (7, 10) | 8 (6, 11) | 7 (5, 9) | 7.5 (7, 9) |
| *Soleus* test units ($\Delta F$) | 3 (2, 4) | 2 (2, 7) | 5 (3, 5) | 3 (2, 4) |
| *Tibialis anterior* motor units | 20.5 (16, 26) | 17 (14, 23) | 17 (13, 23) | 17 (14, 22) |
| *Tibialis anterior* test units ($\Delta F$) | 10 (7, 14) | 10 (6, 10) | 8 (5, 13) | 8 (5, 13) |

**Experiment 2**
| Group | Older adults | Young adults |
| --- | --- | --- |
| *Soleus* motor units | 6 (5, 7) | 5 (4, 8) |
| *Soleus* test units ($\Delta F$) | 3 (1, 4) | 2 (1, 5) |
| *Tibialis anterior* motor units | 19 (15, 26) | 18 (11, 20) |
| *Tibialis anterior* test units ($\Delta F$) | 11 (7, 14) | 14 (5, 17) |

Note: The presented numbers represent the quantity of motor units tracked over time (before and after each condition) in Experiment 1 and tracked between conditions (vibration and control) in Experiment 2.

## Notes

https://github.com/orssatto/PICs-ageing_2.0

### Competing Interest Statement

The authors have declared no competing interest.

